# Group size and social interactions’ strength determine collective response to perturbation in a model of burst-and-coast swimming fish

**DOI:** 10.1101/2025.07.02.662737

**Authors:** Guozheng Lin, Ramón Escobedo, Zhangang Han, Clément Sire, Guy Theraulaz

## Abstract

Collective responses to localized perturbations are essential for the adaptability of animal groups. Using a biologically grounded computational model of burst-and-coast swimming in *Hemigrammus rhodostomus*, we investigate how group size and the strength of social interactions shape collective dynamics under perturbation. The model integrates experimentally derived attraction and alignment rules, behavioral heterogeneity, and boundary effects within a circular tank. We identify four collective states (schooling, milling, turning, and swarming) and characterize a critical regime in which groups exhibit multistable dynamics. At this critical point, small subsets of perturbing individuals, defined by altered social interaction strengths, can induce sharp transitions to new collective states. In particular, such transitions occur in large groups (N = 100) but not in smaller ones (N = 25 or 50), highlighting a size-dependent sensitivity to disturbance. We show that both the nature of the perturbing individuals and the initial state of the group modulate the system’s responsiveness. Our findings suggest that large groups may exploit criticality to remain both robust, flexible, and that individual variability can serve as a catalyst for adaptive reconfiguration. This work provides new insights into how the internal group structure and perturbation design influence collective behavior in animal groups.

## I. INTRODUCTION

Collective motion in fish schools is one of nature’s most fascinating and complex behaviors. Fish exhibit highly coordinated movements that help them evade predators, locate food, and maintain group cohesion [1–3]. The fundamental question that has intrigued researchers for decades is how individual fish interact with their neighbors to produce such synchronized movement patterns [4–6]. Recent advances combining high-resolution tracking in experimental studies with data-driven modeling have shed light on the mechanisms underlying collective motion [7–10]. A key step has been to disentangle and model the respective roles of social interactions (i.e., attraction, alignment, and repulsion), which govern how fish respond to their neighbors while swimming in groups [11, 12]. This made it possible to estimate, for every fish within a school and at every moment, the respective influence that each of its neighbors has on its behavior [13]. Several studies have also shown that to coordinate their movements, fish filter the information coming from their social environment and only pay attention to a small number of neighbors whose identity regularly changes [13–18].

At collective scale, fish schools exhibit various movement patterns in order to adapt to changing environments, including schooling (polarized movement) during foraging, milling (circular motion), swarming (disorganized movement), and directed motion in response to external stimuli (e.g., predators, food sources) [19– 22]. Several studies have demonstrated that the interplay between attraction and alignment forces dictates which state emerges, with phase diagrams mapping transitions between these states [23–26]. Notably, the transition region between milling and schooling appears to be a critical state that optimizes responsiveness to perturbations, suggesting an evolutionary advantage for fish groups that can swiftly switch between these configurations [27]. Some works have recently found signatures of a critical state in fish schools through the analysis of spontaneous behavioral cascades, where large turns in the direction of motion of individuals propagate across a group [28, 29]. Understanding the behavioral mechanisms that drive transitions between these states is essential for predicting school behavior under different environmental conditions.

In a recent work, we showed that schools of rummy-nose tetras (*Hemigrammus rhodostomus*) can tune their collective state when they experience stress [27]. The fish achieve this by adjusting the strength of their social interactions in a way that leads the whole school into a critical state, thus enhancing its sensitivity to perturbations and facilitating rapid adaptation. However, the level of stress of fish is itself modulated by the size of the group, so that a small group operates in a critical state while an additional level of stress is needed to lead a large group of fish in the critical regime.

Therefore, the collective response of a school of fish to a perturbation such as a sudden change in the behavior of a single or a few individuals depends both on group size and on the collective state in which the school finds itself. Moreover, when sudden perturbations occur, social interactions between fish may become asymmetric [30]. In those situations, individuals that first perceive the disturbance reduce their response to neighbors and react preferentially to the perturbing fish. The latter thus exert a disproportionate influence on the rest of the group, which may ultimately trigger a group-wide transition, in a manner analogous to avalanches in critical systems [29, 31, 32]. Even in the absence of external stimuli, the presence of behaviorally heterogeneous individuals—those with heightened sensitivity or exploratory tendencies—can similarly lead to collective transitions [33, 34].

Here, we investigate these mechanisms in greater detail, using a biologically grounded model that allows us to simulate how fish schools respond to localized perturbations under different internal states and group sizes. The model introduced in [11] and [13, 25] describes the interactions arising in burst-and-coast swimming in groups of *H. rhodostomus*. We introduce the behavioral heterogeneity among fish by assigning different attraction and alignment strengths to a small subset of individuals (perturbing individuals), compared to the rest of the homogeneous group (“normal” individuals). Unlike previous work [25], the simulations are conducted in a circular tank to account for boundary effects, which are often present but neglected in experimental and field conditions [7, 35]. We first explore the parameter space of the model applied to groups of *N* = 25, 50, and 100 fish by varying its main parameters, which are the intensity of attraction and alignment, identifying a critical point where multiple states coexist. We then examine how different types of perturbation affect the long-term behavior of a school at the critical point, identifying the interaction parameters of the perturbing fish that have the most significant impact. Finally, we assess how perturbations alter collective behavior when the group starts in a stable schooling or milling state.

## II. METHODS

### A. Model description

We use the model introduced by Calovi *et al*. [11] to describe the burst-and-coast swimming mode of rummy-nose tetra and the social interactions (*Hemigrammus rhodostomus*) when swimming in pairs, later extended by Xue *et al*. [36] to describe collective behavior in different illumination conditions. The swimming pattern of this fish species consists of repeated sequences, each comprising a short and abrupt acceleration, called a “kick”, followed by a quasi-passive deceleration during which the fish glides in a straight line until the next kick.

The model consists of three equations per fish, which determine the instant at which the fish performs a kick and how its position and heading are updated at each kicking instant. At a kicking time 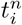, the fish *i* is at position 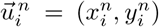 with velocity 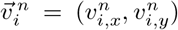, which determines its heading 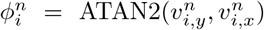 (Figure 1(a)).

**FIG. 1.**
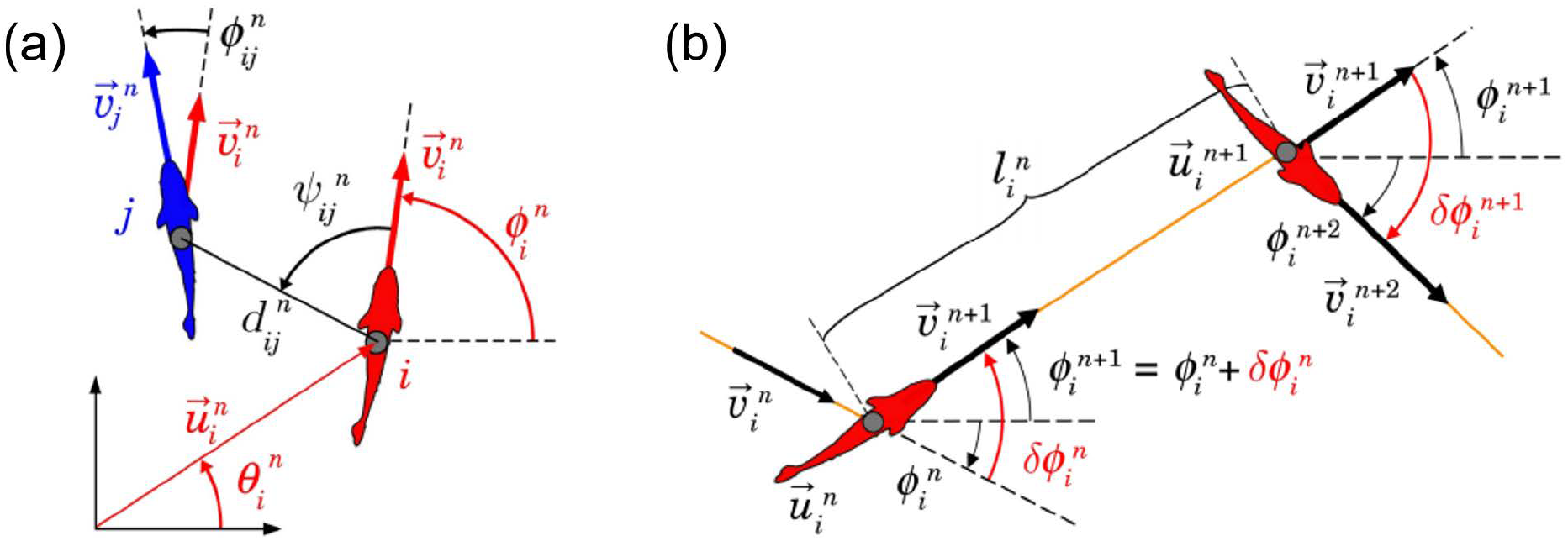
(a) Individual (red) and social (black) state variables of fish *i* with respect to fish *j* (blue) at the instant *t*^*n*^ when fish *I* performs its *n*-th kick: 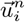 and 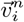 are the position and velocity vectors of fish 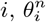 and 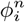 are the angles these vectors make with the horizontal line,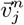 is the velocity vector of fish *j* at time 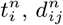 is the distance between fish *i* and 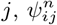 is the angle with which fish *i* perceives *j* (not necessarily equal to 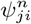), and 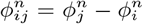 is the heading difference between the two fish. (b) Schematic of the *n*-th kick of fish *i*, moving from 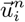 at time 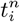 to 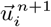 at time 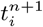 along a distance 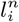. Orange lines denote the fish’s trajectory, black wide arrows denote velocity vectors, and curved arrows represent angles. The heading angle change of fish *i* at time 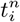 is 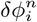. Fish heading during its *n*-th kick is given by 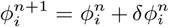. Red angles show the heading variation of fish *i* at the kicking instants 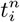 and 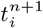.

The *n*-th kick is characterized by its length 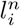 and duration 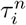. At the end of the kick, the new position 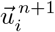 and heading are given by the following equations:

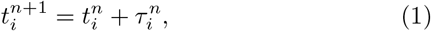

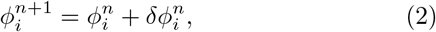

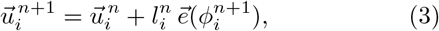

where 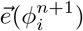 is the unit vector along the angular direction 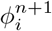, which is the fish heading angle during its *n*-th kick, and 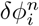 is the heading change from kick *n* − 1 to kick *n*.

The length and duration of a kick performed by a fish are independent of those of its previous kicks, and also of the kicks performed by other fish. When swimming in groups, the kicks of different fish are asynchronous and not necessarily of the same length. Their values are randomly sampled from a bell-shaped distribution,

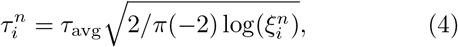

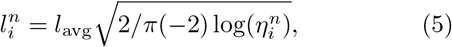

where *τ*_avg_ and *l*_avg_ are the average kick duration and the average kick length, measured experimentally in [36], and 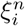 and 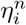 are two independent uniformly distributed random variables between 0 and 1.

According to the experimental observations of Calovi *et al*. [11], the exponential decay of the speed during the gliding phase has a mean relaxation time *τ*_0_, so that the speed at the kicking instant 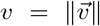 is linked to the kick duration and length by the relation *l* = *vτ*_0_[1 − exp(−*τ/τ*_0_)], allowing us to obtain the position of an individual at any moment during a kick by means of the following expression,

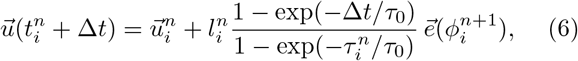

where 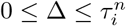.

We assume that the heading angle change of fish *i* results from the additive combination of its spontaneous behavior, the physical constraints of the environment (tank wall and obstacles), and the social interactions with other fish:

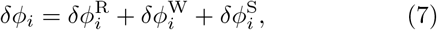

where indices R, W, and S are for Random, Wall, and Social, respectively. Analytical expressions for these interactions were derived from experimental data by means of an extraction procedure in [11] and [36]:

- Heading change due to fish spontaneous behavior *δϕ*^R^:

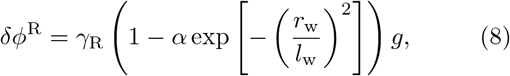

where *γ*_R_ is the *noise* intensity, 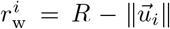 is the distance of fish *i* to the wall, *l*_w_ corresponds to the distance at which the effect of the wall is perceived by the fish, and *g* is a Gaussian random variable with mean 0 and variance 1. We consider that fish spontaneous behavior is conditioned by the vicinity of the wall so that random fluctuations are reduced near the wall (*α <* 1). The numerical values of these parameters were derived from experimental measurements in [36].
- Heading change due to wall repulsion *δϕ*^W^:

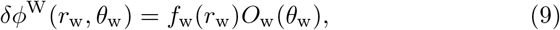

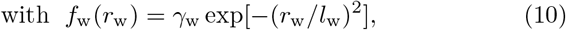

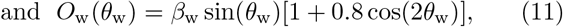

where *f*_w_ is the repulsion strength in function of the distance to the wall *r*_w_, and *O*_w_ is the modulation of this interaction with respect to the angle of incidence to the wall *θ*_w_. The intensity and range of the wall repulsion are given by *γ*_w_ and *l*_w_, and *β*_w_ is a normalization factor.
- Heading change due to social interactions *δϕ*^S^:

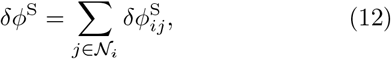

where 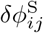 is the pairwise interaction between *i* and *j*, and 𝒩_*i*_ is the set of neighbors interacting with *i. H. rhodostomus* typically interact with two neighbors [13, 16]. These neighbors are selected as the two most influential ones, meaning that their absolute instantaneous contribution to the heading angle change, 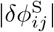, are the two largest ones. This selection reduces the amount of information that needs to be processed in the brain and avoids cognitive overload. As the relative states of fish change with time, the identity of the two most influential neighbors can change accordingly at each kicking instant.

The pairwise interaction function 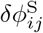 depends on the three variables that determine the relative state of two fish: the distance between them *d*, the angle with which the neighbor is perceived by the focal individual *ψ*, and the difference in heading angle Δ*ϕ* = *ϕ*_*j*_ − *ϕ*_*i*_. Assuming no side preference in fish, a symmetry constraint applies, 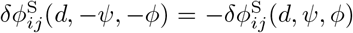. This allows us to separate the function into two contributions, attraction and alignment [*]. Finally, we assume variables separability. In summary [12, 36] :

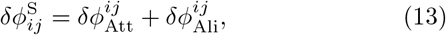

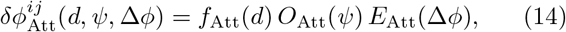

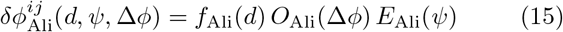

where

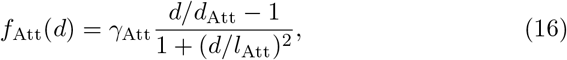

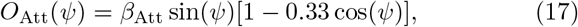

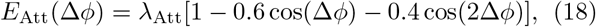

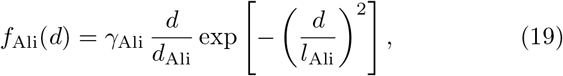

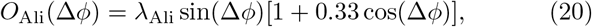

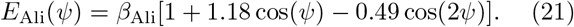

The parameters *γ*_Att_ and *γ*_Ali_ represent the strength of the attraction and alignment interactions, respectively. The parameters *l*_Att_ and *l*_Ali_ define the respective range of these interactions, while *d*_Att_ and *d*_Ali_ are characteristic distances: at *d*_Att_, attraction and repulsion balance each other, whereas *d*_Ali_ simply makes *f*_Ali_ dimensionless. The constants *β*_Att_, *β*_Ali_, *λ*_Att_, and *λ*_Ali_ are normalization factors of the angular functions, allowing for direct comparison between them.

We consider that *γ*_Att_ and *γ*_Ali_ are the control parameters that determine the individual behavior of the fish and their collective state in a schooling, milling or swarming formation, while the other parameters, whose values were experimentally obtained in [36], remain constant (Table I).

**TABLE I.** Parameter values used in the model, obtained from [36].

| Parameter | Value | Description |
| --- | --- | --- |
| $l_{\text{avg}}$ | 0.06 | Mean kick length (m) |
| $\tau_{\text{avg}}$ | 0.55 | Mean kick duration (s) |
| $\tau_0$ | 0.87 | Relaxation time during gliding (s) |
| $\gamma_{\text{R}}$ | 0.1 | Noise intensity |
| $\alpha$ | 0.67 | Noise modulation close to the wall |
| $\gamma_{\text{w}}$ | 0.4 | Wall interaction strength |
| $l_{\text{w}}$ | 0.053 | Wall interaction range (m) |
| $\beta_{\text{w}}$ | 1.9612 | Normalization of angular function |
| $l_{\text{Att}}$ | 0.170 | Attraction range (m) |
| $d_{\text{Att}}$ | 0.015 | Attraction balance distance (m) |
| $\beta_{\text{Att}}$ | 1.43 | Normalization of angular function |
| $\lambda_{\text{Att}}$ | 0.9 | Normalization of angular function |
| $l_{\text{Ali}}$ | 0.145 | Alignment range (m) |
| $d_{\text{Ali}}$ | 0.015 | Normalization distance (m) |
| $\beta_{\text{Ali}}$ | 0.9 | Normalization of angular function |
| $\lambda_{\text{Ali}}$ | 1.43 | Normalization of angular function |

### B. Quantification of collective behaviors

The quantification of collective behaviors involves measuring emergent patterns in groups of individuals, with a focus on how local interactions give rise to global collective dynamics. To characterize these behaviors, three key metrics are considered: dispersion, polarization, and milling index.

These metrics are based on the barycenter of the group, whose position and velocity vectors, denoted as 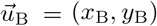 and 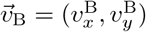, are given by:

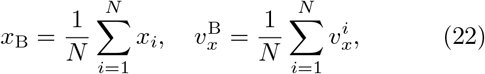

with analogous expressions for the *y*-components. The heading angle of the group barycenter is defined by the orientation of the velocity vector:

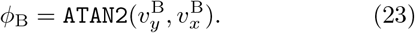

We then define the following collective measures:

#### Group Dispersion

The dispersion of the group, denoted as *D*(*t*) *∈* [0, 1], quantifies the spatial spread of individuals around the group’s barycenter. This measure is normalized by the radius of the circular tank, R, and is defined as:

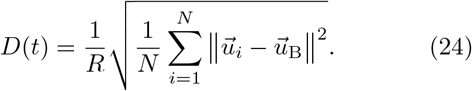

A low value of *D*(*t*) indicates that individuals are closely clustered around the group’s barycenter, reflecting a highly cohesive or compact configuration. Conversely, higher values of *D*(*t*) suggest that individuals are more widely dispersed, indicating a loosely structured or scattered group.

#### Group Polarization

The group polarization, denoted as *P*(*t*) *∈* [0, 1], quantifies the degree of alignment among individuals within the group. It is defined as the norm of the average unit direction vectors, where each individual’s direction is given by 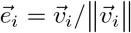:

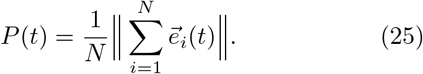

A polarization value close to 1 indicates that most individuals are moving in the same direction, reflecting a highly ordered state. For randomly distributed orientations, *P*(*t*) is of order 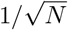, while values near zero occur only when individual orientations perfectly cancel each other out, corresponding to a highly specific and balanced configuration.

#### Milling Index

The milling index, denoted as *M*(*t*) *∈* [0, 1], quantifies the extent to which individuals exhibit rotational movement around the group’s barycenter. It is defined as:

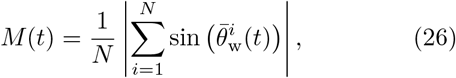

where 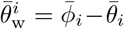 represents the angle of incidence of the velocity vector of individual *i* relative to a circular trajectory centered on the barycenter, within the barycentric reference frame.

In this reference frame, denoted with an overbar, the coordinates and velocity components are given by:

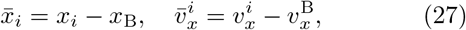

with the angular components defined as:

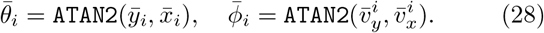

A high value of *M*(*t*) indicates a well-structured rotational movement around the barycenter, characteristic of milling behavior. Conversely, low values suggest the absence of such organized circular motion, indicating a more dispersed or unstructured collective state.

### C. Simulation setup

We conducted long-term simulations of the model across a wide range of attraction and alignment strength parameters, *γ*_Att_ and *γ*_Ali_, to assess the impact of a small fraction of perturbing fish on the collective behavior of the group. The simulations considered a group of *N* fish swimming in a circular tank of radius *R*, with a subset of *N*_p_ individuals classified as perturbing fish. These perturbing fish were assigned distinct attraction and alignment strength values, 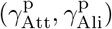, while the “normal” fish used the parameters 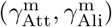. All other model parameters were identical for both perturbing and normal fish and remained constant throughout the simulations.

To investigate the effects of group size and the fraction of perturbing fish, we have considered three different group sizes: *N* = 25, 50, and 100. The proportion of perturbing fish (PPF) was defined as PPF = *N*_p_*/N* and varied between 0% and 16%. To maintain a consistent group density across different sizes, the radius of the circular tank was set to 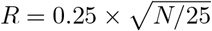 for each scenario.

For all combinations of group size (*N* = 25, 50, 100) and PPF, each simulation was run for 1500 seconds, starting from a randomly initialized state in terms of position and heading. Only the final 600 seconds of each simulation, after the system reached behavioral stability, were used for statistical analysis. Each parameter combination was replicated 10 times for control conditions (without perturbing fish) and 5 times for experimental conditions (with perturbing fish), ensuring robustness in the results.

## III. RESULTS

### A. Collective states without perturbations

We first analyze the behavior of fish schools in the absence of perturbing individuals as a baseline for comparison. To establish this benchmark, we observe the long-term dynamics of fish schools under different parameter combinations of 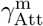 and 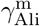 across all group sizes. Our simulations reveal four distinct collective states: schooling, milling, turning, and swarming (see Supplementary Material (SM) Movie S1-S4). These states are not easily distinguishable using a single state variable, such as *P* or *M* alone, but can be effectively characterized by their combined values [24, 37]. Figure 2(a)-(d) presents snapshots of 100 fish exhibiting each of these states, alongside the probability density distributions (PDF) of *P* and *M* for the corresponding simulations. In the schooling state, fish align and move in the same direction (see SM Movie S1), leading to higher values of *P* and lower values of *M*. In the milling state, fish collectively rotate around a central point (see SM Movie S2), resulting in lower values of *P* and higher values of *M*. In the turning state, fish exhibit coordinated directional changes (see SM Movie S3), with both *P* and *M* concentrated at higher values. Finally, in the swarming state, fish movement is relatively disordered (see SM Movie S4), yielding low values for both *P* and *M*.

**FIG. 2.**
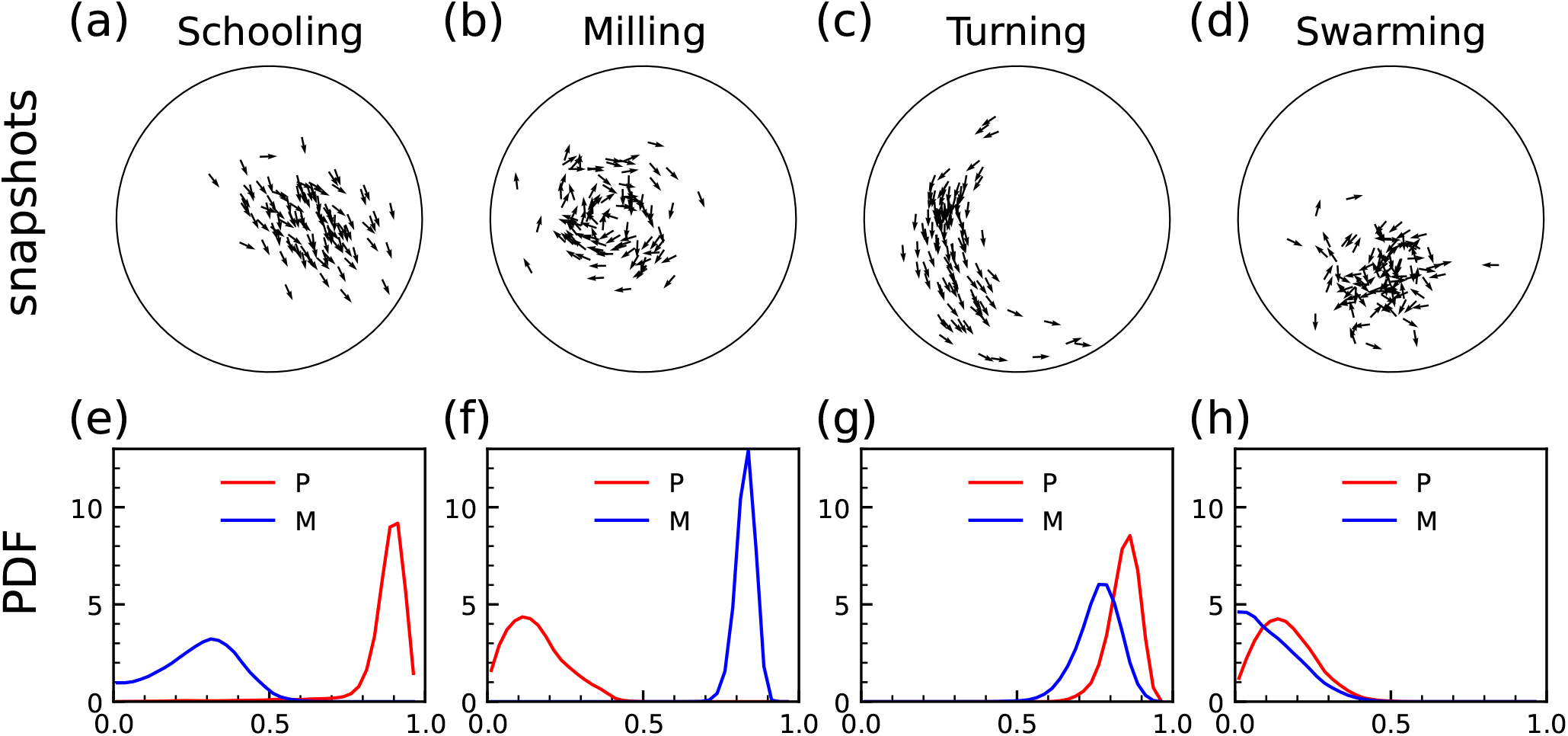
Snapshots (first row) and probability density functions (PDF) of polarization and milling index during motion (second row) of four typical collective states at *N* = 100. (a)(e) Schooling, where the fish swim in the same direction, corresponding to large *P* and small *M*. (b)(f) Milling, the fish rotate around a certain center, corresponding to large *M* and small *P*. (c)(g) Turning, the fish show an extended curved pattern with a tendency to turn in a certain direction, corresponding to large *P* and large *M*. (d)(h) Swarming, the fish move in a disorganized direction, corresponding to small *P* and small *M*.

Figure 3 illustrates the phase diagrams of fish schools in the parameter space 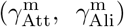 for three different group sizes: *N* = 25, 50, and 100. To facilitate state classification, we categorize the values of *P* and *M* into the four observed states using a threshold of 0.5, as indicated in the color map to the right of the figure: schooling occurs when *P >* 0.5 and *M <* 0.5; milling when *P <* 0.5 and *M >* 0.5; turning when *P >* 0.5 and *M >* 0.5; and swarming when *P <* 0.5 and *M <* 0.5.

**FIG. 3.**
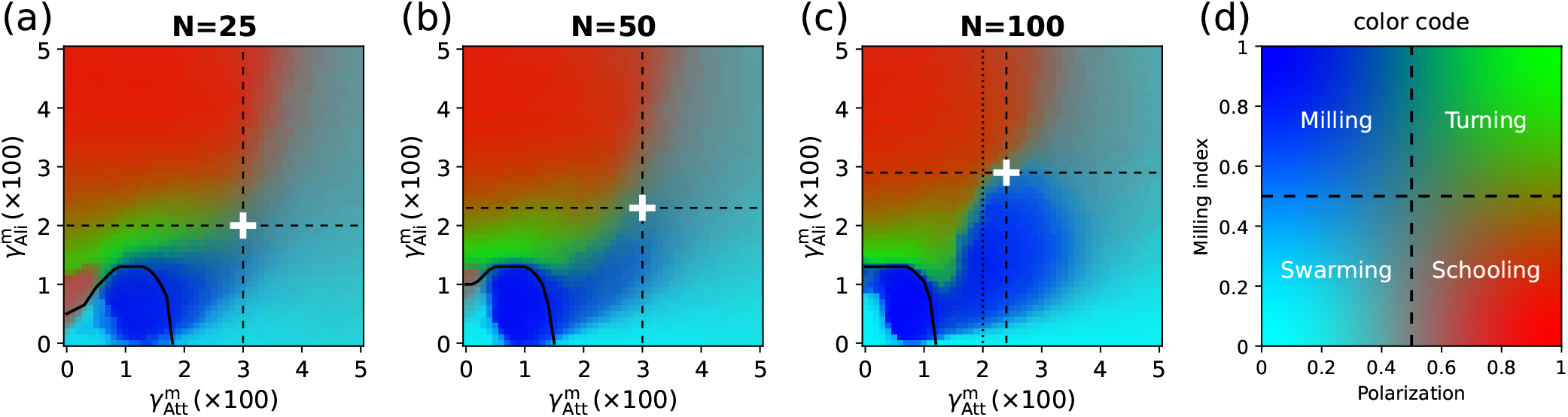
Phase diagrams of fish schools in the parameter space 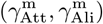 for different group sizes. The right panel displays the corresponding color map, where each (*P, M*) pair is assigned a specific color value, thereby distinguishing the four states: Schooling, Milling, Turning, and Swarming. The white crosses indicate the intersection points of all four states, corresponding to the critical point where both *P* and *M* are equal to 0.5. The parameter values for critical points are: (0.03, 0.02) for *N* = 25, (0.03, 0.023) for *N* = 50 and (0.024, 0.029) for *N* = 100. The black curve in the lower-left corner of each panel represents the threshold where dispersion is 0.65, with regions below the curve exhibiting dispersion values greater than 0.65.

Our analysis indicates that all four states emerge across different group sizes. When the attraction strength is higher than 0.035, fish schools predominantly exhibit the swarming state. Conversely, when 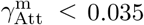, the region corresponding to the milling state expands with increasing group size. Notably, the milling behavior observed in a circular tank can be classified into two distinct types: (i) an intrinsic milling motion that arises independently of the boundary (see SM Movie S2) and (ii) a boundary-induced milling motion, where the group rotates along the tank walls (see SM Movie S5). These two types can be distinguished based on the dispersion *D* of the group, with the former exhibiting a higher degree of aggregation and the latter displaying more dispersion. In Figure S1, the time series of two milling states are presented in detail. The milling index values of the intrinsic milling motion and the boundary-induced milling motion are both stable above 0.75, but the corresponding dispersions are significantly different, with the former always remains below 0.4, while the latter always remains above 0.65. In the lower-left region of the phase diagram, where both attraction and alignment strengths are low, we plot a black curve indicating the threshold of average dispersion at *D* = 0.65. Below this curve, where dispersion exceeds 0.65, weak fish-fish interactions are still sufficient to induce a milling-like state due to boundary constraints, whereas milling states found elsewhere arise intrinsically from collective interactions.

An intriguing feature of the phase diagram is the intersection of all four states at a single point (marked by a white cross in the diagram), occurring in the region where fish schools remain sufficiently aggregated (above the black curve, where dispersion is below 0.65). At this point, both *P* and *M* are approximately equal to 0.5. Observations reveal that fish schools in this state exhibit multi-stability, frequently transitioning between schooling and milling, with occasional shifts into turning and swarming (as shown in the first 1500 seconds in Figure 4(a)-(c)). This phenomenon bears a striking resemblance to the concept of criticality in statistical physics, where a system at a critical state between macroscopic phases exhibits heightened sensitivity to external perturbations. In such states, minor local perturbations can induce significant system-wide transitions. This concept has been widely applied to biological collective systems, including fish schools and bird flocks, to explain their ability to rapidly respond to dynamic and potentially hazardous environments [38, 39]. Consequently, in the following analysis, we define the white cross-marked point as the *critical point* to assess the response of fish schools to perturbations in this state.

**FIG. 4.**
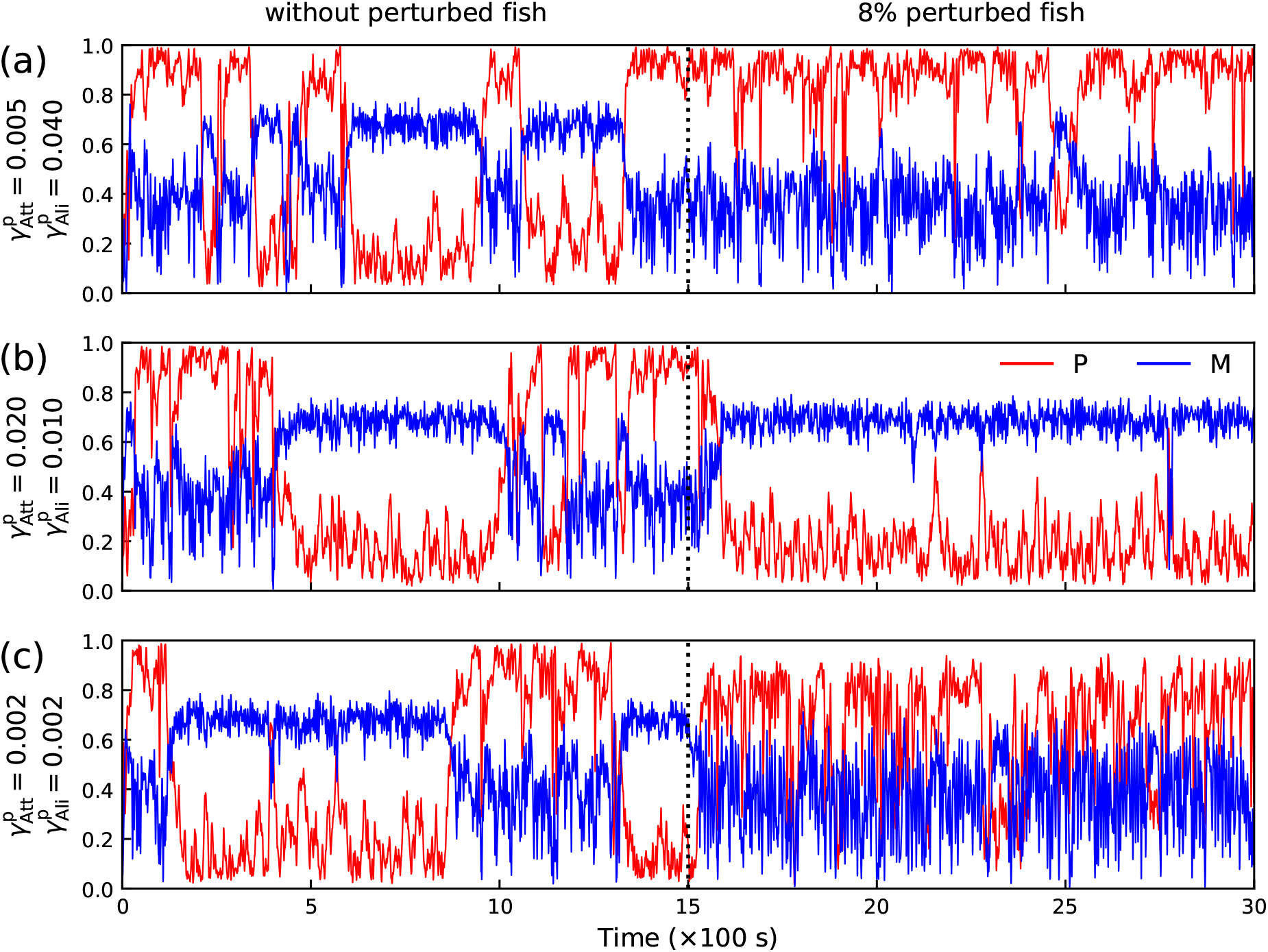
Polarization and milling time series of *N* = 100 fish schools, where the fish school is in a critical state for the first 1500 seconds, and at 1500 seconds 8% of them become perturbing fish, with the parameters of the perturbing fish being (a) 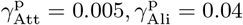, (b) 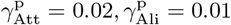 (c) 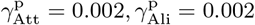.

### B. Collective response to perturbations when a school is in the critical state

In the critical state, defined as the point where the system’s attraction and alignment strengths coincide with the critical point marked by the white cross in Figure 3, small perturbations can cause the system to transition into other collective states. To investigate this phenomenon, we systematically explored the parameter space of perturbing fish, 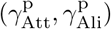, and tested various proportions of perturbing individuals within the group.

Figure 5 illustrates the changes in the system’s state under different combinations of attraction and alignment strengths for perturbing fish. The state transition is quantified by the average difference in polarization and milling index between conditions with and without perturbing fish at the critical point. The first, second, and third columns of the figure correspond to group sizes of 25, 50, and 100, respectively. Notably, in this figure, the proportion of perturbing fish is set to PPF = 8%. Additional simulations with PPF = 4% are presented in Figure S2 in the SM, where the conclusions remain qualitatively unchanged.

**FIG. 5.**
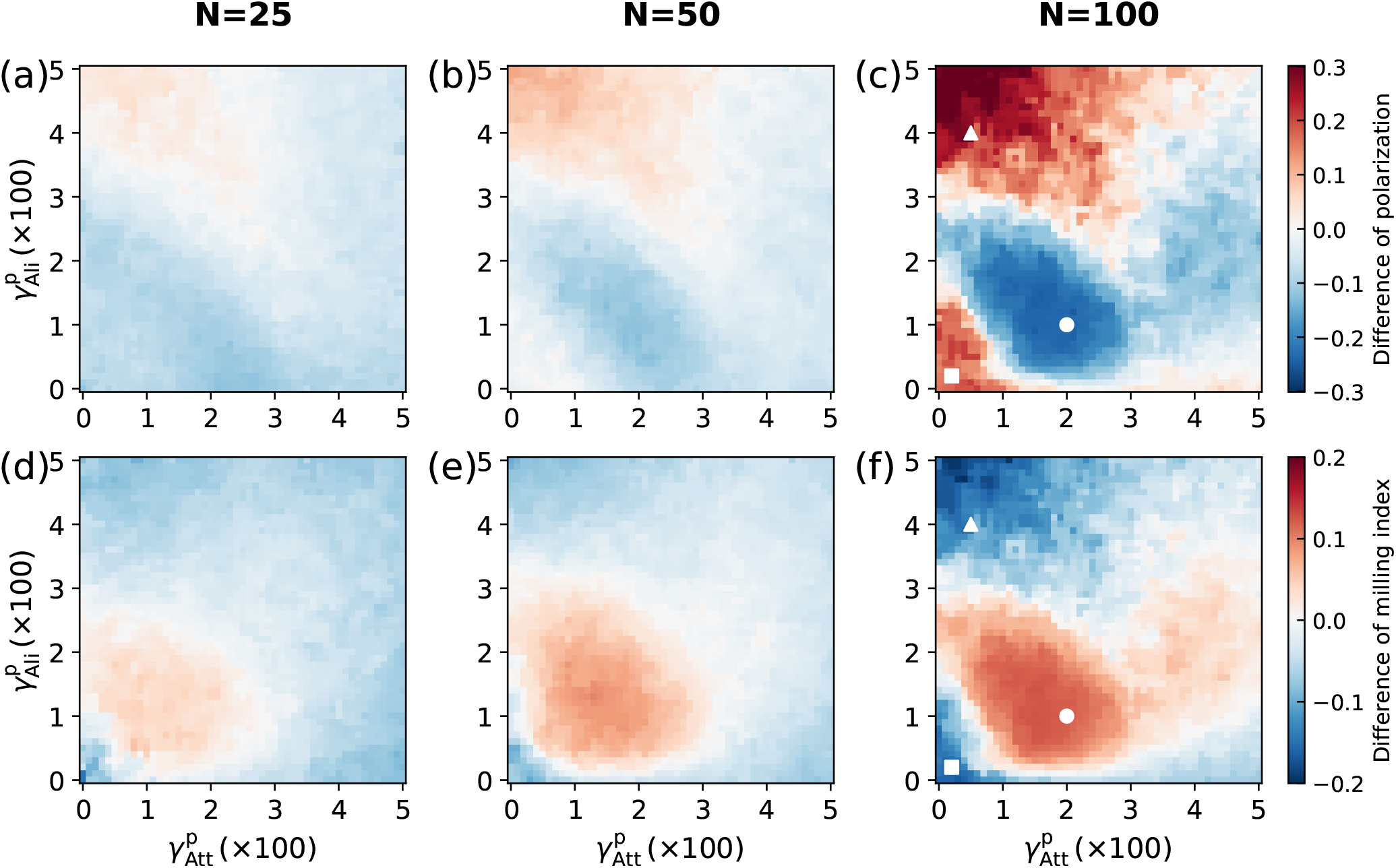
Difference in group polarization and milling index across the 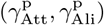 parameter space following the introduction of 8% perturbing fish, compared to the unperturbed case. Each column represents a different group size. The parameters for normal (unperturbed) fish are set at the critical point, indicated by the white cross in Figure 3. White markers highlight regions where significant changes in the collective state of the fish school were observed.

As shown in Figure 5, for group sizes of *N* = 25 and *N* = 50, the variations in polarization and milling index remain relatively small across the parameter space, with absolute differences remaining below 0.1. This suggests that a small proportion of perturbing fish has little impact on the overall collective state. However, for a group size of *N* = 100, distinct regions emerge where the absolute difference exceeds 0.2, indicating that 8% of perturbing fish can induce substantial shifts in the collective state of the school. Furthermore, as shown in Figure S1, even a PPF of 4% is sufficient to cause significant changes in polarization or milling index when *N* = 100. These results show that group size strongly influences its collective responses to disturbances affecting the behavior of a small proportion of individuals when the group is in a critical state.

The effects of perturbing fish on the *N* = 100 group are particularly pronounced in three parameter regions, as marked by the white symbols in Figures 5(c) and (f): (i) Near the triangular marker 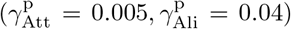, where attraction strength is low and alignment strength is high, perturbation leads to a significant increase in group polarization. (ii) Near the circular marker 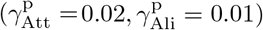, where attraction strength is moderate and alignment strength is low, perturbation results in a substantial increase in the milling index. (iii) Near the square marker 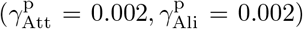, where both attraction and alignment strengths are weak, perturbation causes a slight increase in polarization.

Although perturbing fish induce measurable changes in the mean polarization and milling index, this alone does not confirm whether the school transitions from a multistable state at the critical point to a single stable state. To address this, we analyzed the time series (Figure 4) and probability density functions (PDFs, Figure 6) of polarization and milling index for the three white-marked parameter points in Figure 5(c)(f), comparing them with the unperturbed case.

**FIG. 6.**
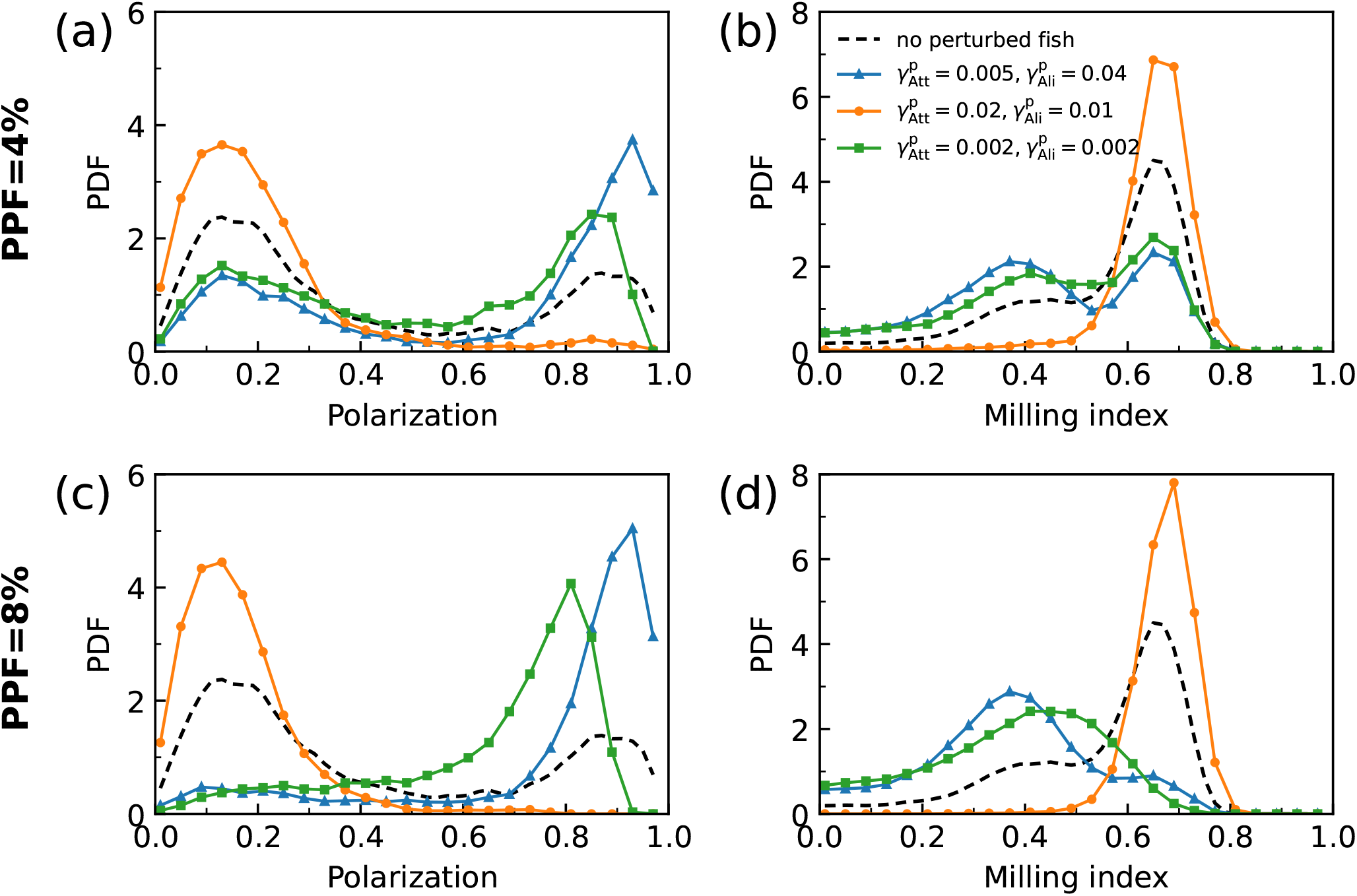
Probability density functions (PDFs) of polarization and milling index for fish schools without perturbing individuals (dashed lines) and after the introduction of perturbing fish (solid lines). The proportion of perturbing fish (PPF) is 4% in the first row and 8% in the second row. The three colored solid lines represent the distributions corresponding to the three sets of perturbing fish parameters, which are associated with the white-marked points in Figure 5. The group size is fixed at *N* = 100, with the parameters of normal (unperturbed) fish set at the critical point.

In Figure 4, we let 100 fish run for 1500 seconds in the critical state, then introduced 8% of perturbing fish and run for another 1500 seconds. In the critical state without perturbing fish, the polarization and milling index of the fish schools change abruptly between high and low values and can remain there for a period of time after the change (as shown in the first 1500 seconds, see also the first 30s of SM Movie S6-S8), causing the fish schools to jump between different stable states. At this point, a small number of perturbing fish can direct them to jump to one of these states (as shown in the last 1500 seconds, see also the last 30s of SM Movie S6-S8). This was not the case for 25 and 50 fish (see Figures S3 and S4 in SM), where the fish schools also showed frequent shifts between high and low values of polarization and milling index, but the shifts did not show persistence in the stable state. This suggests that the collective response produced by 100 fish is fundamentally different from that of smaller 25 or 50 fish, with the former showing discontinuous transitions in bistable states and the latter continuous transitions in different states.

Figure 6 illustrates more clearly the effect of perturbing fish on the state of N = 100 fish school. In the absence of perturbing fish, the group polarization exhibits a clear bimodal distribution, with both high and low values, while the milling index also shows a bimodal tendency, albeit less pronounced—indicating system instability at the critical point. However, when perturbing fish with strong alignment strengths are introduced (blue curves with triangles), the polarization distribution shifts toward higher values, and the milling index shifts toward lower values. As PPF increases from 4% to 8%, the bimodal distributions gradually transition to uni-modal, signifying a shift from the critical state to a stable schooling state. When perturbing fish with moderate attraction and weak alignment strengths are introduced (orange curves with circles), the polarization distribution concentrates at lower values while the milling index peaks at higher values, forming a single-peaked distribution regardless of PPF, indicating a transition to a stable milling state. Finally, when perturbing fish with both weak attraction and weak alignment strengths are introduced (green curves with squares), the polarization distribution shifts toward higher values, while the milling index remains concentrated below 0.5. This concentration effect becomes more pronounced as PPF increases, suggesting a transition from the critical state to the schooling state.

The effect of different numbers of perturbing fish on *N* = 100 fish schools is further shown in Figure 7. When perturbing fish with weak attraction and strong alignment strengths are introduced, the polarization increases approximately linearly with the number of perturbing fish, and the milling index decreases approximately linearly. Note that, like in physics, the slope of these linear variations can be associated to a susceptibility, measuring the sensitivity to the perturbation [30]. Again, in analogy with the fluctuation-dissipation theorem in physics, the susceptibility has been shown to be proportional to the variance/fluctuations of the corresponding order parameter (polarization or milling) [30].

**FIG. 7.**
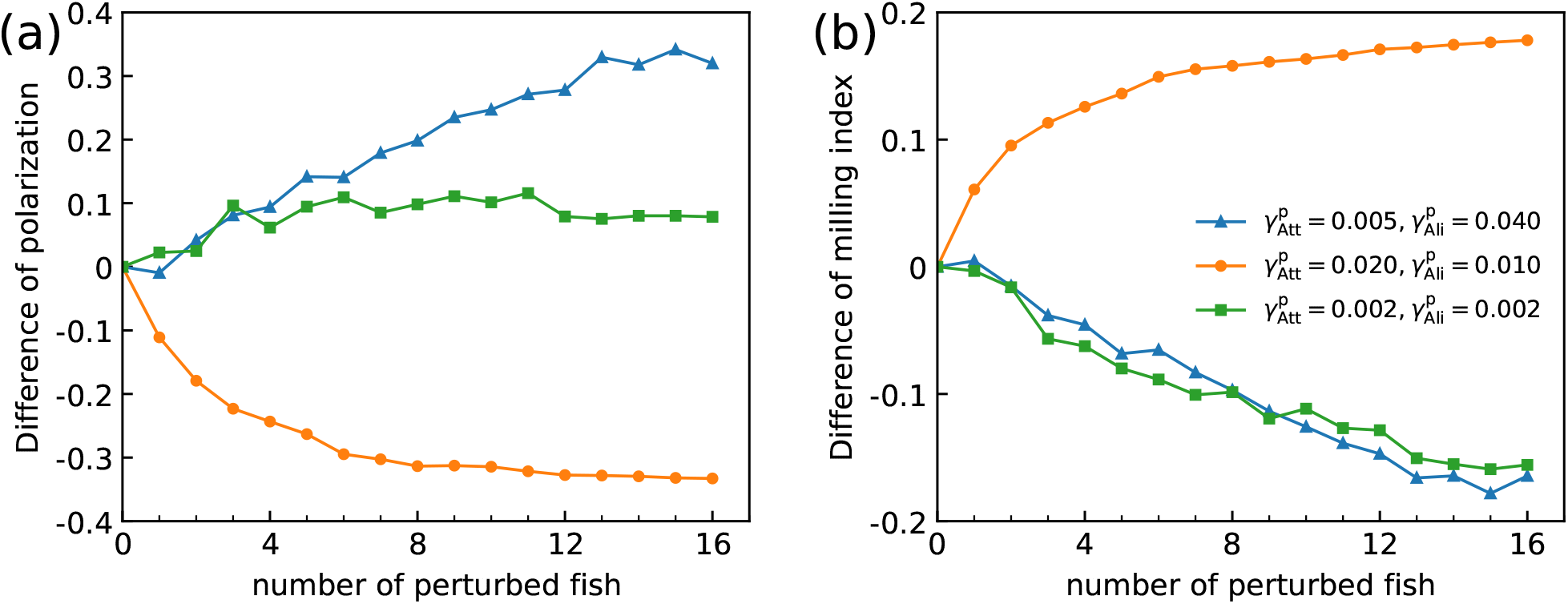
Variation of the difference in polarization and the difference in milling index as a function of the number of perturbing fish. The group size is fixed at *N* = 100, the parameters for normal fish are fixed at the critical point 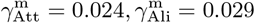.

When perturbing fish with moderate attraction and weak alignment strengths are introduced, an increase in the number of perturbing fish caused a rapid decrease in polarization and a rapid increase in milling at the very beginning (number ¡ 8), but continued addition of perturbing fish thereafter would not continue to enhance the effect (number ¿ 8). When perturbing fish with both weak attraction and weak alignment strengths are introduced, as the number of perturbing fish increases, polarization consistently increases by about 0.1, while milling index decreases approximately linearly.

### C. Collective response to perturbations when a school is in the schooling or milling state

In this section, we examine the system’s response to perturbations when it is in a more stable schooling or milling state, rather than in the critical state. As previously observed, for groups of 25 and 50 fish, even when the system is in a critical state, a small proportion of perturbing individuals has little effect on the group’s collective state. Therefore, we focus our analysis on the group of 100 fish.

In Figure 3(c), we highlight the straight line 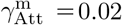, along which the system transitions from the milling state to the schooling state as the alignment strength 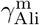 increases. Based on this observation, we set the parameters of the normal fish to vary along this straight line while assigning the perturbing fish parameters to the values corresponding to the white markers in Figure 5(c), which represent three parameter regions where perturbations exert a significant influence.

Figure 8 illustrates the effect of perturbing fish on the group polarization (first row) and milling index (second row) as the alignment strength of the normal fish, 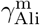, varies, while the attraction strength remains constant at 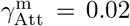. Each column represents a different set of parameter values for the perturbing fish.

**FIG. 8.**
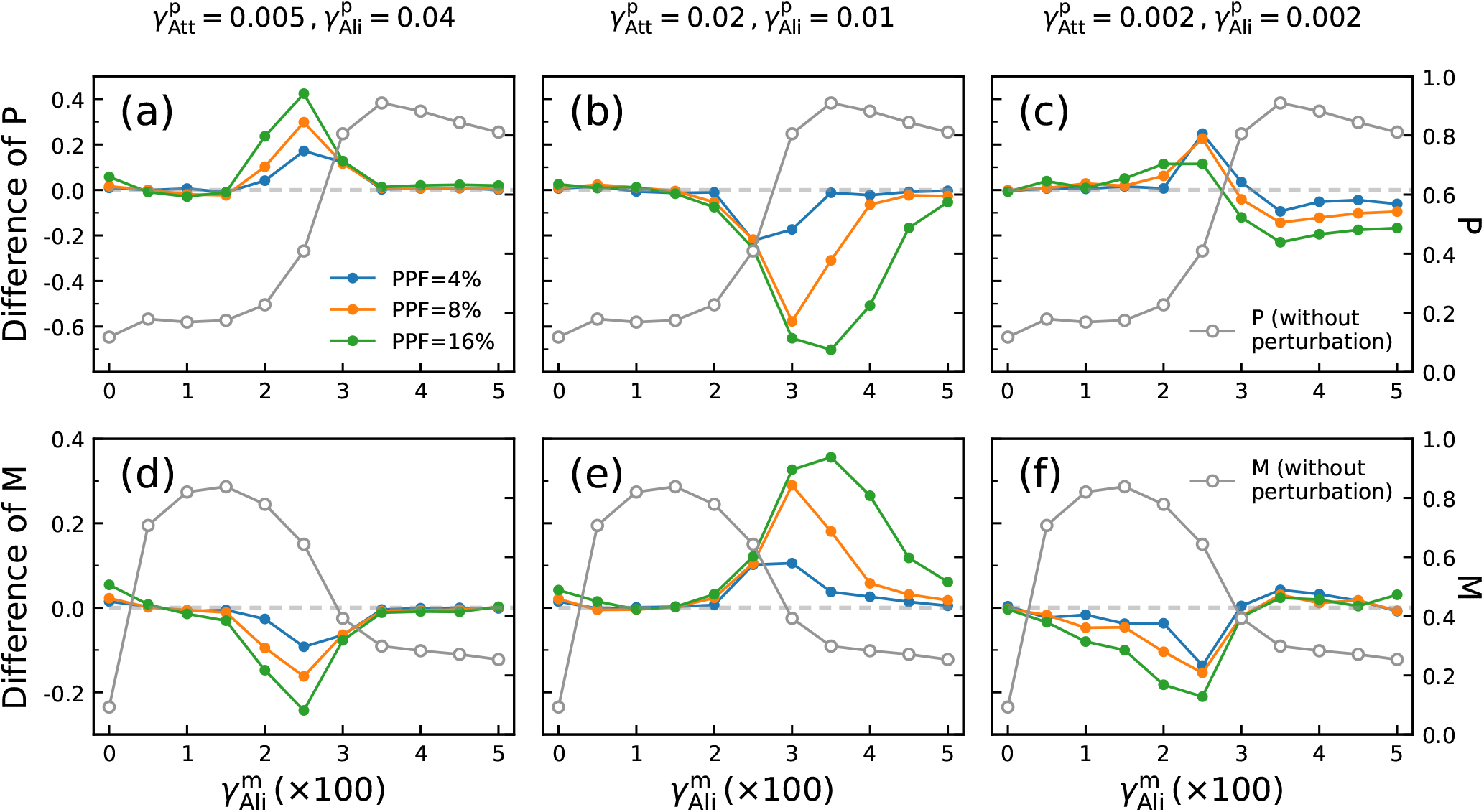
Variation of difference of polarization and milling index as a function of 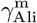. Each column corresponds to a different set of perturbing fish parameters, while each curve represents a different proportion of perturbing fish. As a comparison, the variation of polarization and milling index with 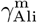 in the absence of perturbation are displayed on the right-side axis, represented by the solid gray line with hollow circles. The gray dashed lines are the zero axis of the left y-axis.

In the absence of perturbing fish (solid gray lines with hollow circles, right-side y-axis), polarization gradually increases while the milling index decreases as 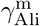 increases from 0.005, leading to a transition from a stable milling state 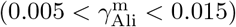 to a stable schooling state 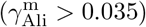.

When perturbing fish with stronger alignment strength are introduced (Figures 8(a) and (d)), their presence has little impact on the system when it is in a highly polarized schooling state 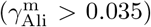 or a strongly milling state 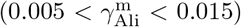. However, in the transition region 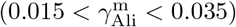, introduction of perturbing fish causes an increase in polarization and a decrease in milling index, and the magnitude of the effect increases further as PPF increases. This indicates that perturbing fish with higher alignment strength facilitate the transition from the milling state to the schooling state.

When perturbing fish with moderate attraction strength and low alignment strength are introduced (Figures 8(b) and (e)), they have little effect when the system is in a stable milling state 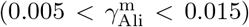. However, as the system evolves toward schooling 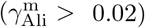, an increasing number of perturbing fish leads to a sharp drop in polarization and a corresponding rise in the milling index. This suggests that perturbing fish with these social interaction parameters promote a transition from the schooling state back to the milling state.

When perturbing fish with both weak attraction and weak alignment strengths are introduced (Figures 8(c) and (f)), a natural consequence is observed: these socially weaker individuals disrupt the overall order of the fish school, reducing both high polarization values and high milling index values. However, an interesting effect emerges when the system is in an intermediate state, transitioning from low to high polarization (i.e.,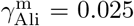). In this case, the introduction of perturbing fish unexpectedly increases the system’s polarization from below 0.5 to above 0.5, while simultaneously decreasing the milling index. In fact, this is consistent with Figure 7, where individuals with weak attraction and alignment strengths can still increase the polarization of the group in the critical state. This finding suggests that perturbing fish with weak social influence do not merely disrupt collective behavior but can, under specific conditions—such as the intermediate state between milling and schooling—enhance the overall order of the group.

## IV. DISCUSSION

How local perturbations affect the collective behavior is a crucial question for understanding the adaptability of complex living systems. In this article, we have explored the collective response of fish schools when only a few individuals behave differently, using a biologically grounded computational burst-and-coast swimming model. We introduced a small fraction of perturbing fish, i.e., individuals with different social interaction strength, into otherwise uniform schools. By running simulations in a circular tank, we have investigated how the group’s collective state changes in response to these perturbations. We further examined how factors such as group size and initial collective state affect a school’s ability to shift behavior due to a few perturbing individuals.

In the absence of perturbing fish, the results of our simulations confirm those obtained with other models of collective swimming involving other forms of social interactions and continuous movement [24, 30, 37, 40]. Fish schools can display four distinct collective states in phase space: schooling, milling, turning, and swarming. As shown in Figure 3, the region corresponding to the milling state expands with increasing group size. This trend is consistent with the results of the burst-and-coast model presented in [25], which does not account for boundary effects. However, in contrast to [25], the boundary-inclusive model used here still displays a well-defined milling state in the lower-left region of the phase space, corresponding to low attraction and alignment strengths. Indeed, both modeling and experimental studies [35, 40] have emphasized that state transitions in fish schools are fundamentally altered by the presence of boundaries. When boundaries are taken into account, bistability can be dissipated, leading the group to transition from schooling or swarming into a milling state.

The model further indicates that the nature of the critical state depends on group size. Large fish schools (*N ≥* 100) exhibit abrupt transitions between distinct stable states (as seen in the first 1500 seconds of Figure 4), whereas smaller schools (*N* < 100) transition more gradually between states (see the first 1500 seconds in Figures S3 and S4). This suggests that larger schools are more sensitive to perturbations, with even small disturbances capable of triggering rapid switches between stable collective behaviors. This scaling effect becomes even more apparent when perturbing individuals are introduced into the simulation. In small groups of 25 or 50 fish, behavioral changes remained minimal even when up to 8% of individuals were perturbing fish (Figure 5). In contrast, in large schools of 100 fish, the same proportion of perturbed individuals frequently induced abrupt and persistent transitions to different collective states (Figure 5). These results are fully consistent with the phenomenology of critical systems in physics, where the susceptibility/sensitivity to a perturbation is proportional to the fluctuations of the order parameter without perturbation, and both diverge at a critical point as the size *N* of the system goes to infinity (thermodynamic limit). This scaling effect highlights how group size can fundamentally alter a system’s sensitivity to disturbance. It also suggests that fish may form large schools not just for safety in numbers, but also to increase behavioral flexibility. Larger groups appear more capable of quickly reorganizing in response to minimal but well-targeted disruptions, thus facilitating their foraging for food or responding to environmental stimuli [28, 41]. This challenges the idea that group robustness is purely about averaging out noise [42]. Instead, large groups may be more adaptable precisely because they can be nudged toward new states more easily when needed [43].

The model also reveals that the type of behavioral change in the group depends on the nature of the perturbing fish. If the perturbing individuals have high alignment but low attraction, they tend to lead the group toward a schooling state (blue lines in Figure 7). If their attraction is moderate and alignment is low, the school shifts toward a milling state (orange lines in Figure 7). Interestingly, even fish with both weak alignment and attraction, which might be expected to cause disorder, can sometimes help stabilize a group by nudging it toward a more cohesive state. These effects are strongest when the school is large and near the transition region (green lines in Figure 7 and Figure 8(c, f)), again, consistent with the physics of critical phenomena [30]. When the group is in a transition state, these individuals that weakly interact with the other fish actually increase group order. One possible explanation is that these weakly social fish are less influenced by other fish and thus are more likely to move in a straight line, thus guiding other fish in a consistent direction.

We also used the model to test what happens when perturbing fish are added to groups already in a stable schooling or milling state. In these conditions, the effects are more limited, but still informative. Perturbing fish with high alignment help reinforce schooling behavior and can even trigger transitions from milling to schooling. Conversely, fish with moderate attraction and weak alignment can pull a group out of a schooling state and back into milling. These findings show that group behavior is not just passively resistant to change in stable states. Depending on the context, perturbing individuals with different social interaction rules can either stabilize or destabilize group patterns. Experimental studies have shown that less social or “bold” individuals are more likely to assume leadership roles within animal groups, often guiding their conspecifics away from danger or toward novel food sources [34]. Similar dynamics have been observed in bird flocks, where changes in the alignment and attraction behaviors of perturbing individuals can lead to rapid collective maneuvers, such as flock-wide swerving to avoid hazards [44, 45]. In all of these contexts, diversity in behavior might thus benefit group decision-making or adaptive responses.

In summary, this study highlights how interaction strength and group size shape the dynamics of collective behavior. More importantly, our results indicate that perturbations are not always disruptive. Depending on their nature, they can enhance collective order or guide the group to a more adaptive configuration.

## ACKNOWLEDGMENTS

GL was supported by the Fundamental Research Funds for the Central Universities (2024XKRC078) and a grant from the China Scholarship Council (CSC N°202006040162). ZH was supported by the National Natural Science Foundation of China (Grant No. 62176022). GT, RE and CS were supported by the Agence Nationale de la Recherche (ANR-20-CE45-0006-1). RE was also supported by the Spanish State Research Agency (AEI, Ministerio de Ciencia, Innovación y Universidades) through project PID2020-115088RBI00/AEI/10.13039/501100011033

