## Supplementary Information for "Group size and social interactions’ strength determine collective response to perturbation in a model of burst-and-coast swimming fish" (from another version of this preprint: version 1, 2025-07-03)

### (a) Supplementary figures

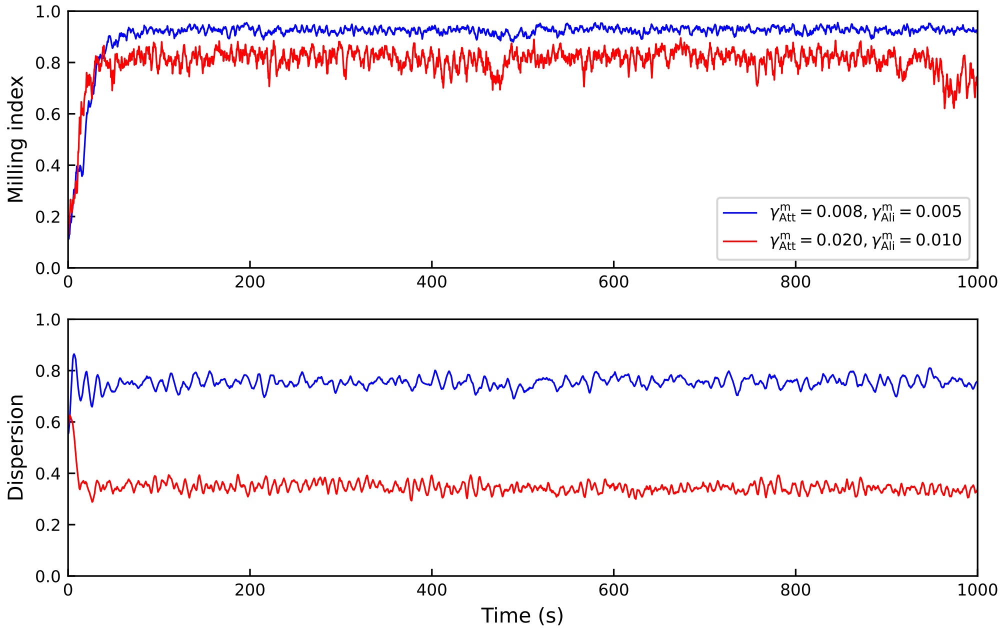


**Figure S1.** Times series of milling index and dispersion of the two kind of milling motion. The intrinsic milling motion corresponding to red curves, with parameters $(\gamma^{\mathrm{m}}_{\mathrm{Att}}, \gamma^{\mathrm{m}}_{\mathrm{Ali}}) = (0.02, 0.01)$, and boundary-induced milling motion corresponding to blue curves, with parameters $(\gamma^{\mathrm{m}}_{\mathrm{Att}}, \gamma^{\mathrm{m}}_{\mathrm{Ali}}) = (0.008, 0.005)$.

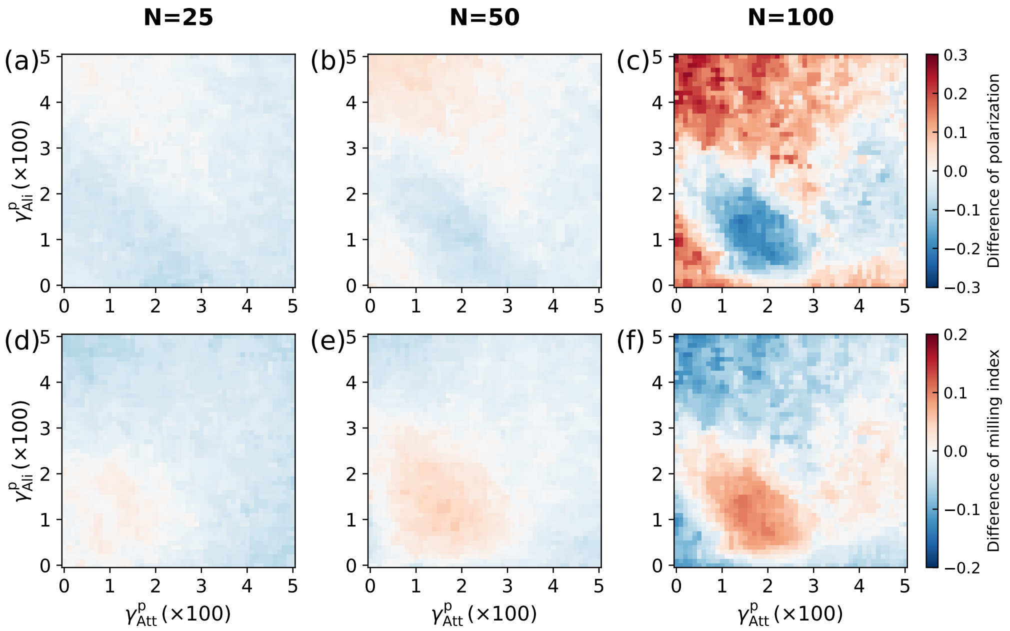


**Figure S2.** Difference in group polarization and milling index across the $(\gamma^{\mathrm{P}}_{\mathrm{Att}}, \gamma^{\mathrm{P}}_{\mathrm{Ali}})$ parameter space following the introduction of 4% perturbed fish, compared to the unperturbed case. Each column represents a different group size.

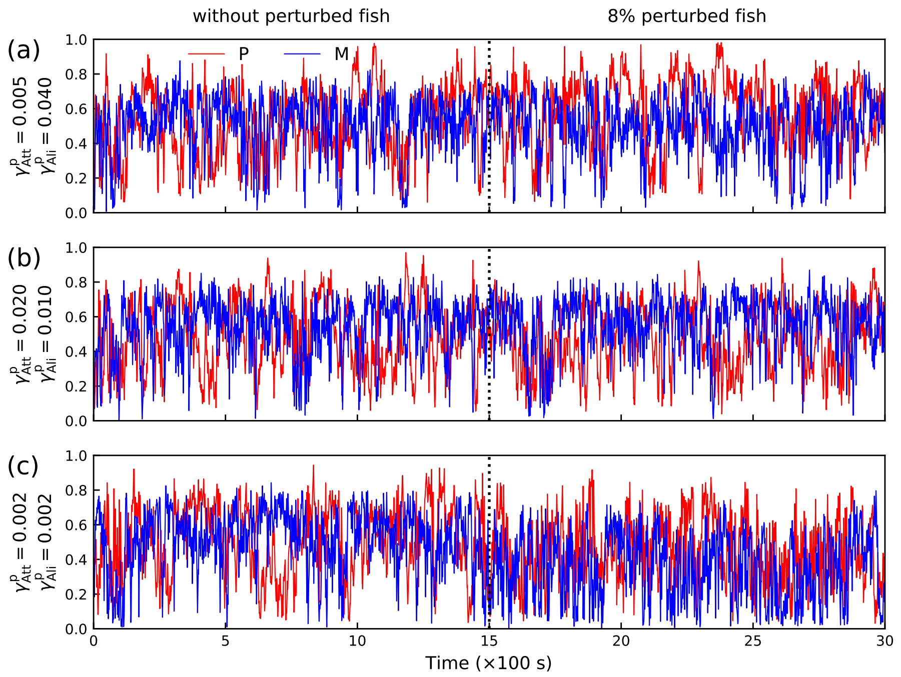


**Figure S3.** Polarization and milling time series of $N = 25$ fish schools, where the fish school is in a critical state for the first 1500 seconds, and at 1500 seconds 8% of them become perturbed fish, with the parameters of the perturbed fish being (a) $\gamma^{\mathrm{P}}_{\mathrm{Att}} = 0.005$, $\gamma^{\mathrm{P}}_{\mathrm{Ali}} = 0.04$, (b) $\gamma^{\mathrm{P}}_{\mathrm{Att}} = 0.02$, $\gamma^{\mathrm{P}}_{\mathrm{Ali}} = 0.01$, (c) $\gamma^{\mathrm{P}}_{\mathrm{Att}} = 0.002$, $\gamma^{\mathrm{P}}_{\mathrm{Ali}} = 0.002$.

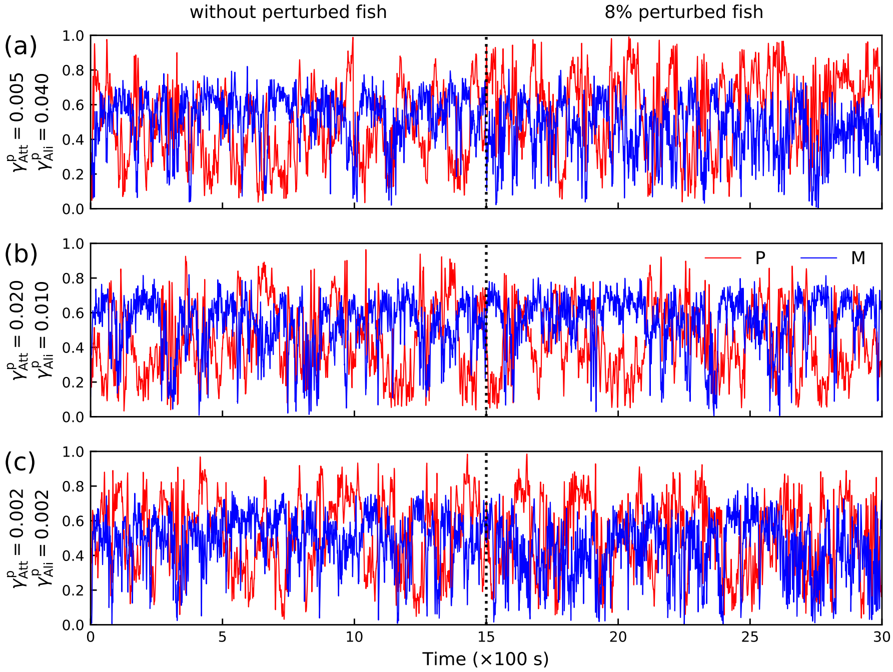


**Figure S4.** Polarization and milling time series of $N = 50$ fish schools, where the fish school is in a critical state for the first 1500 seconds, and at 1500 seconds 8% of them become perturbed fish, with the parameters of the perturbed fish being (a) $\gamma^{\mathrm{P}}_{\mathrm{Att}} = 0.005, \gamma^{\mathrm{P}}_{\mathrm{Ali}} = 0.04$, (b) $\gamma^{\mathrm{P}}_{\mathrm{Att}} = 0.02, \gamma^{\mathrm{P}}_{\mathrm{Ali}} = 0.01$, (c) $\gamma^{\mathrm{P}}_{\mathrm{Att}} = 0.002, \gamma^{\mathrm{P}}_{\mathrm{Ali}} = 0.002$.

### (b) Supplementary videos





- Movie S1: **Schooling state**. Video excerpt of a simulation with a group of 100 fish swimming in a circular tank of radius 50 cm with $(\gamma^{\mathrm{m}}_{\mathrm{Att}}, \gamma^{\mathrm{m}}_{\mathrm{Ali}}) = (0.01, 0.04)$ and without perturbing fish. Each point represents current position of each fish and the tail of the point represents the trajectory within 0.4 seconds. The video has been speed up by 4 times.
- Movie S2: **Milling state**. Video excerpt of a simulation with a group of 100 fish swimming in a circular tank of radius 50 cm with $(\gamma^{\mathrm{m}}_{\mathrm{Att}}, \gamma^{\mathrm{m}}_{\mathrm{Ali}}) = (0.02, 0.01)$ and without perturbing fish. Each point represents current position of each fish and the tail of the point represents the trajectory within 0.4 seconds. The video has been speed up by 4 times.
- Movie S3: **Turning state**. Video excerpt of a simulation with a group of 100 fish swimming in a circular tank of radius 50 cm with $(\gamma^{\mathrm{m}}_{\mathrm{Att}}, \gamma^{\mathrm{m}}_{\mathrm{Ali}}) = (0.01, 0.015)$ and without perturbing fish. Each point represents current position of each fish and the tail of the point represents the trajectory within 0.4 seconds. The video has been speed up by 4 times.
- Movie S4: **Swarming state**. Video excerpt of a simulation with a group of 100 fish swimming in a circular tank of radius 50 cm with $(\gamma^{\mathrm{m}}_{\mathrm{Att}}, \gamma^{\mathrm{m}}_{\mathrm{Ali}}) = (0.03, 0.002)$ and without perturbing fish. Each point represents current position of each fish and the tail of the point represents the trajectory within 0.4 seconds. The video has been speed up by 4 times.
- Movie S5: **Boundary-induced milling state**. Video excerpt of a simulation with a group of 100 fish swimming in a circular tank of radius 50 cm with $(\gamma^{\mathrm{m}}_{\mathrm{Att}}, \gamma^{\mathrm{m}}_{\mathrm{Ali}}) = (0.008, 0.005)$ and without perturbing fish. Each point represents current position of each fish and the tail of the point represents the trajectory within 0.4 seconds. The video has been speed up by 4 times.
- Movie S6: **Introducing perturbing fish with** $(\gamma^{\mathrm{p}}_{\mathrm{Att}}, \gamma^{\mathrm{p}}_{\mathrm{Ali}}) = (0.005, 0.04)$ **at critical state**. In the first 30 seconds of the movie, fish group is in a critical state with parameters $(\gamma^{\mathrm{m}}_{\mathrm{Att}}, \gamma^{\mathrm{m}}_{\mathrm{Ali}}) = (0.024, 0.029)$, and then 8% of them transforms into perturbing fish (marked in red color) with parameters $(\gamma^{\mathrm{m}}_{\mathrm{Att}}, \gamma^{\mathrm{m}}_{\mathrm{Ali}}) = (0.005, 0.04)$. Each point represents current position of each fish and the tail of the point represents the trajectory within 0.4 seconds. The video has been speed up by 4 times. The video has been speed up by 10 times.
- Movie S7: **Introducing perturbing fish with** $(\gamma^{\mathrm{p}}_{\mathrm{Att}}, \gamma^{\mathrm{p}}_{\mathrm{Ali}}) = (0.02, 0.01)$ **at critical state**. In the first 30 seconds of the movie, fish group is in a critical state with parameters $(\gamma^{\mathrm{m}}_{\mathrm{Att}}, \gamma^{\mathrm{m}}_{\mathrm{Ali}}) = (0.024, 0.029)$, and then 8% of them transforms into perturbing fish (marked in red color) with parameters $(\gamma^{\mathrm{m}}_{\mathrm{Att}}, \gamma^{\mathrm{m}}_{\mathrm{Ali}}) = (0.02, 0.01)$. Each point represents the current position of each fish and the tail of the point represents the trajectory within 0.4 seconds. The video has been speed up by 4 times. The video has been speed up by 10 times.
- Movie S8: **Introducing perturbing fish with** $(\gamma^{\mathrm{p}}_{\mathrm{Att}}, \gamma^{\mathrm{p}}_{\mathrm{Ali}}) = (0.002, 0.002)$ **at critical state**. In the first 30 seconds of the movie, fish group is in a critical state with parameters $(\gamma^{\mathrm{m}}_{\mathrm{Att}}, \gamma^{\mathrm{m}}_{\mathrm{Ali}}) = (0.024, 0.029)$, and then 8% of them transforms into perturbing fish (marked in red color) with parameters $(\gamma^{\mathrm{m}}_{\mathrm{Att}}, \gamma^{\mathrm{m}}_{\mathrm{Ali}}) = (0.002, 0.002)$. Each point represents the current position of each fish and the tail of the point represents the trajectory within 0.4 seconds. The video has been speed up by 4 times. The video has been speed up by 10 times.
